# Performance of the BD Phoenix CPO detect assay for the detection and classification of OXA-48 producing-*Escherichia coli* that do not coproduce ESBL/p*AmpC*

**DOI:** 10.1101/2024.04.29.591738

**Authors:** Fátima Galán-Sánchez, Inés Portillo-Calderón, Manuel Rodriguez-Iglesias, Álvaro Pascual, Lorena López-Cerero

## Abstract

Reliable detection of OXA-48-like enzymes in routine antimicrobial susceptibility testing is difficult, especially if the isolate does not coproduce other beta-lactamases. The BD Phoenix™ Emerge panel includes the CPO Detect Test (CPO-T), which allows simultaneous antimicrobial susceptibility testing and carbapenemase detection and characterization. We evaluated the accuracy of detection and classification of carbapenemases determined by CPO-T against a characterized set of OXA-48-producing *Escherichia coli* isolates that do not co-produce ESBL/pAmpC, and BD Phoenix™ for rapid antimicrobial susceptibility testing. A total of 51 *E. coli* isolates were included All the isolates were sequenced using Illumina and analyzed using Resfinder and CARD for resistance determinants, MLSTfinder for typing and Clermont Phylotyper for phylotypes. CPO-T was performed within the BD Phoenix™ NMIC-502 panel. Disk diffusion, with readings at 4, 6, 8 and 24 hours, was performed as reference method (RDD). The CPO-T detected carbapenemase activity in 100% of the isolates within 7 hours of incubation and 42 (82.4%) isolates were correctly assigned to the class D Ambler. For all isolates and antibiotics, BD Phoenix™ provided definitive susceptibility results in less than 8 hours in 64.6% of cases compared to 58.2% with RDD. Overall categorical agreement was 90.7% and 5.9% very major, 6.2% major and 3.2% minor errors were observed. Our results demonstrate that CPO-T is a reliable tool for the detection of OXA-48 in *E. coli* isolates that do not co-produce ESBL/p*AmpC* without the need for additional testing, and that BD Phoenix™ provides rapid results for the antibiotics most commonly used in empirical treatment.

**IMPORTANCE:** Detection of OXA-48-like enzymes in isolates that do not co-produce other beta-lactamases in routine antimicrobial susceptibility testing is difficult, and studies on these types of isolates are scarce. Our study analyzes the performance of the BD Phoenix™ Emerge panel, which includes the CPO detection test (CPO-T), in a characterized set of OXA-48-producing *Escherichia coli* isolates that do not co-produce ESBL/pAmpC, and also provide information about time to results of this system.

## INTRODUCTION

OXA-48-type β-lactamases are increasingly reported in enterobacterial species. In a recent multicenter study conducted in Spain (1), *bla*^OXA–48^ was predominant (73.1%) in carbapenemase-producing *Escherichia coli*. These enzymes hydrolyze penicillins, but not broad-spectrum cephalosporins, and have an attenuated ability to hydrolyze carbapenems. This weak carbapenem hydrolytic activity can lead to false carbapenem susceptibility testing results. For this reason, reliable detection of OXA-48-like enzymes usually requires immunochromatographic or molecular methods, as biochemical assays like Carba-NP are less reliable than they are for metallobetalactamases or KPC enzymes (2).

This characteristic weak-hydrolysis activity of OXA-48-type β-lactamases (3) can complicate its detection in routine antimicrobial susceptibility testing, especially if the isolate does not coproduce other types of β-lactamases. Recent studies have demonstrated the ability of BD Phoenix™ CPO Detect test (CPO-T) to detect carbapenemases, using phenotypic or molecular detection assays as reference methods (4-9). These studies found similar high overall sensitivities of 89.4% - 100% for the detection of carbapenemase activity in different collections of clinical *Enterobacterale*s isolates. In these studies, CPO-T sensitivity for OXA-48-like enzyme detection was 97.1% - 100%, although the isolates collections tested contained most OXA-48 producers with other enzymes like ESBL or pAMPC.

Recently, several promising new antimicrobial susceptibility tests systems have been developed to provide rapid testing (rAST), within 8h or less, but these new technologies will also require optimizations and are usually costly (10). Rapid reading with routinely used traditional methods such as disk diffusion have been developed and validated for blood cultures (https://www.eucast.org/rapid_ast_in_bloodcultures). Some conventional automated systems, such as BD Phoenix™, could provide rapid susceptibility results for some antibiotics, as it uses both an oxidation-reduction indicator and turbidity growth detection.

We evaluated the accuracy of CPO-T for the detection and classification of carbapenemase production of a characterized set of OXA-48 producing *E. coli* isolates that do not co-produce ESBL/p*AmpC*, and the time to results of the BD Phoenix™ system was compared to disk diffusion, including reading at shortened incubation times.

## MATERIAL AND METHODS

### Collection of isolates

A total of 51 OXA-48-producing and ESBL/p*AmpC*-negative *E. coli* isolates were submitted to the Andalucía Reference laboratory during the period 2018-2022. Andalusian microbiology laboratories included in the PIRASOA program voluntarily submitted carbapenemase-producing Gram negative bacteria, detected according to EUCAST guidelines for the detection of resistance mechanisms, which include a screening cut-off of ertapenem > 0.125 mg/l for *Enterobacteriaceae*. (https://www.eucast.org/fileadmin/src/media/PDFs/EUCAST_files/Resistance_mechanisms/EUCAST_detection_of_resistance_mechanisms_170711.pdf).

The isolates were detected from 40 (78,4%) surveillance cultures and 11 (21,6%) clinical samples (5 blood, 4 exudate, 1 respiratory and 1 urine samples). They were collected from 10 hospitals located in 5 Andalusian provinces.

Initial bacterial identification was confirmed using the MALDI-TOF Biotyper (Bruker Daltonics). Antimicrobial susceptibility testing was performed using the commercial MDRM1 panel (Microscan; Beckman Coulter), which includes ertapenem concentrations of 0.12, 0.5 and 1 mg/l, and susceptibility disk diffusion on Mueller Hinton agar (Oxoid) for ertapenem, imipenem, meropenem, ceftazidime and cefotaxime, following EUCAST guidelines. Phenotypic carbapenemase characterization was performed with the imipenem hydrolysis test (β-CARBA test, Bio-Rad), immunochromatography with the NG-test CARBA 5 (NG Biotech) and study of inhibitors with disk diffusion (dipicolinic acid, phenyl-boronic acid, and cloxacillin). In addition, all the isolates were sequenced with Illumina MiSeq 2000 by using Nextera Flex DNA library preparation and reads were submitted to Enterobase and assembled de novo with EToKi (Enterobase Tool Kit). Quality parameters are included in Table S1. Assemblies were annotated by using Resfinder (11) with the default configuration (60-90% threshold) for acquired resistance determinants, PointFinder (12) and CARD (13) for SNPs variants within *gyrA/parC* topoisomerases genes, and regulatory network of AcrAB efflux pump (*acrR, marAR* and *soxS/R*), MLSTfinder for typing and Clermont Phylotyper to assign phylotype.

### BD Phoenix™ CPO detect test

The test was performed within the BD Phoenix™ NMIC-502 panel (BD Diagnostic Systems, Sparks, MD) according to the manufacturer’s’ recommendations. Briefly, the panels were manually inoculated with isolates prepared using samples stored at -80°C storage, after an overnight growth onto BD Columbia Agar with 5% Sheep Blood. The BD Phoenix™ NMIC-502 routine panel includes a confirmatory test that utilizes meropenem, doripenem, temocillin and cloxacillin, alone and in combination with various chelators and beta-lactamase inhibitors in amounts required for the detection and classification of carbapenemase-producing organisms (CPO-T), along with 136 wells containing antimicrobial agents for antimicrobial susceptibility testing. Results were interpreted by BD Epicenter software (BD Diagnostic Systems, Sparks, MD). This software allows reporting of results for each antibiotic at an individual level in real time, i.e., as soon as the result was finalized. Isolates were considered to be susceptible according to EUCAST S and I breakpoints.

### Disk diffusion test and readings (RDD)

Susceptibility disk diffusion was performed on Mueller Hinton agar (BD) for all antibiotics included in the NMIC-502 panel, following EUCAST guidelines, but with readings of inhibition zones at 4, 6, 8 and 24 hours of incubation. Inhibition zones were only read when zone edges were clearly visible, and for β-lactams, an inhibition of 100% was considered at the end of incubation. For both, the BD Phoenix and RDD methods, QC strains (*E*.*coli* ATCC 25922) were included. Readings at 24 hours were considered as the reference method, and at the point of 8 hours ± 5 minutes were used to provide data on the time to results. A Chi-squared test was used to compare the results for BD Phoenix and RDD at 8 hours ± 5 minutes (considered significant if p<0.05). ATU results for ciprofloxacin were not included for minor errors (mE) calculation and were only arose when comparing susceptible and resistant BD Phoenix to intermediate RDD results.

Rates of categorical agreement (CA), mE, major errors (ME), and very major errors (VME) were defined according to the US Food and Drug Administration (FDA guidelines) (14). As recommended by FDA, congruent expected performances were CA >90%, VME <1.5%, and ME < 3%.

## RESULTS

Forty-eight isolates (94.2%) carried *bla*_OXA-48_, two isolates (3.9%) carried *bla*_OXA-244_ and one isolate (1.9%), *bla*_OXA-181._ Additionally, sixteen isolates co-carried *bla*_TEM-1_, one co-carried *bla*_SHV-1_ and another one, *bla*_OXA1_. Twenty-four isolates (47%) carried genes encoding aminoglycoside-modifying enzymes, 21 (41.2%) isolates had genes linked to trimethoprim and/or sulfonamides resistance and 19 (37.2%) isolates had genes linked to tetracyclines resistance (Figure S1).

Quinolone resistance-determining regions (QRDR) mutations and/or plasmid-mediated quinolones resistance were detected in 16 (31.4%) isolates. Several isolates showed mutations: 14 (27,5%) isolates showed at least one mutation in topoisomerases genes (Figure S1), 6 (11,8%) isolates had a variant in the *uhpT* gene (E350Q), 7(13,7%) isolates a variant in the *cyaA* gene (S352T) and 46 (86,3%) had the same SNPs variant in the *marR* gene (Y137H, G103S). Phylogroup B1 (39%) predominated, followed by phylogroup A (21.6%) (Figure S1). The collection was polyclonal and isolates belonged to 36 different MLSTs and 19 (37%) isolates were a member of 3 clonal complexes (10, 23 and 155).

The CPO-T detected carbapenemase production in 100% of the isolates within 7 hours of incubation. The assignments following detection of carbapenemase production by CPO-T were as follows: 42 (82.4%) isolates were assigned to class D Ambler, 5 (9.8%) isolates to class B Ambler, and in 4 (7.8%) were detected as carbapenemase producers without specific assigment. The nine misclassified or unclassified isolates were resistant to ertapenem and piperacillin-tazobactam and susceptible to aztreonam, ceftazidime-avibactam, and meropenem.

Antimicrobial susceptibility results for all the isolates using BD Phoenix™ and disk diffusion are presented in Table 1, along with CA and VME, ME and mE when the BD Phoenix results were compared to disk diffusion with readings at 24 h as reference method for each antibiotic. The percentages of VME exceeded those recommended for cefalexin, cefixime, cefuroxime, ciprofloxacin, cotrimoxazole, imipenem, fosfomycin, tigecycline and gentamycin (Table 1). Overall CA was 90.7%, which is above the recommended threshold.

**Table 1.**
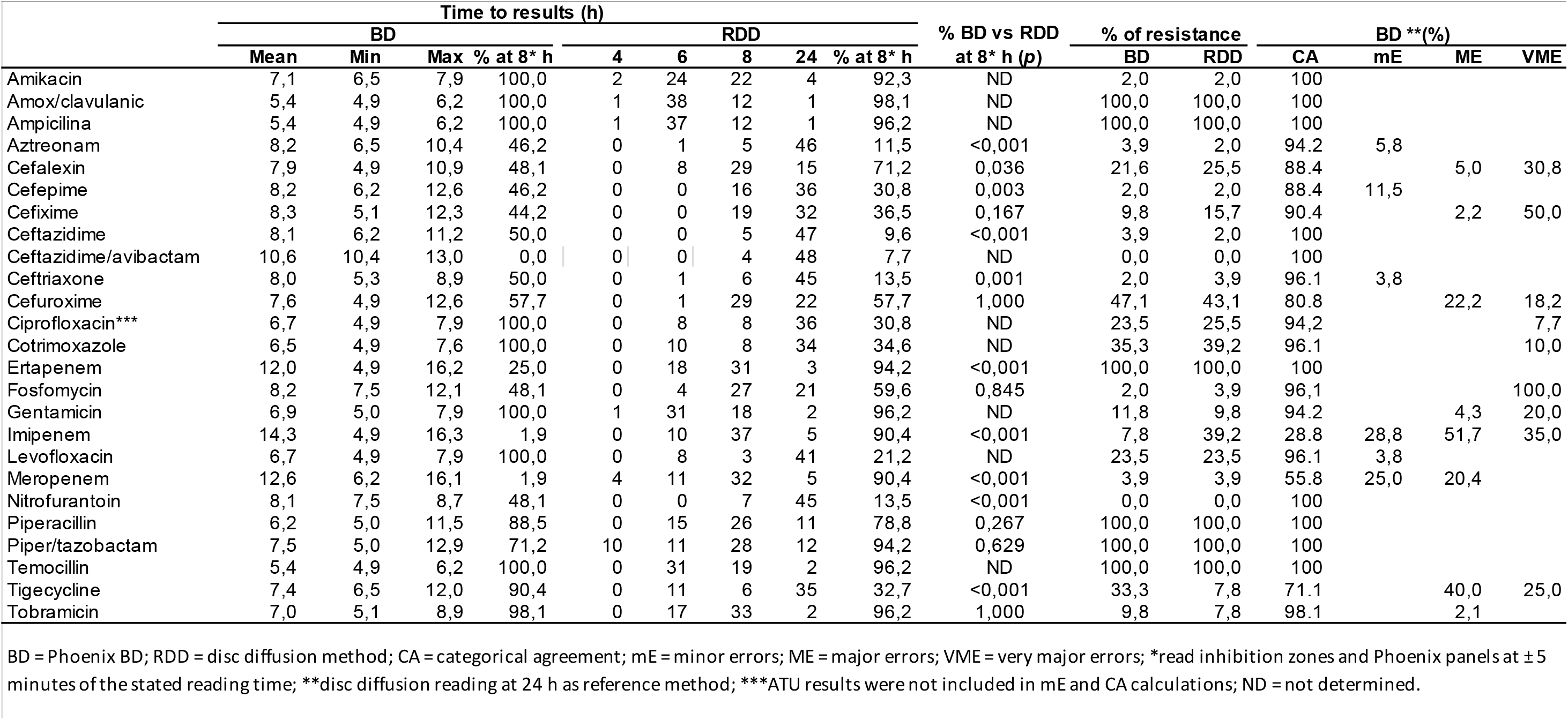
Performance of Phoenix panel compared with disk diffusion method with 51 OXA-48-producing *E. coli* isolates

Imipenem and meropenem showed the lowest CA due to the growth of colonies within the carbapenems inhibition zones. If these two carbapenems are not included, the CA increases to 95.0%. Upon further analysis of the VMEs in the case of cephalosporins, it was observed that none of the isolates resistant by RDD and susceptible by Phoenix BD had mutations in *acrR* and *soxS/R*. Only the following determinants were found in the resistant isolates with cephalosporins VME: *bla*_TEM-1_ in 2 of 4 cephalexin-resistant isolates, and the H45R point mutation in the *AmpC* promoter in 2 isolates (1 resistant to cefixime and 1 resistant to cefixime and cefuroxime). In one isolate, which was found to be resistant by RDD and susceptible by Phoenix to ciprofloxacin, no mutations in topoisomerases or plasmid determinants of quinolone resistance were detected. Similarly, no mutations in the *acrR* or *soxS/R* or *tet* genes were detected in an isolate resistant by RDD and susceptible by BD Phoenix to tigecycline. On the other hand, 2 isolates that gave VME with cotrimoxazole had only *dfrA* genes and no *sul* genes.

Cumulative and average time to result for each antibiotic in the BD Phoenix™ NMIC-502 panel are given in Table 1. For all isolates and antibiotics, BD Phoenix™ provided definitive susceptibility results in less than 8 hours in 64.6% of the results (95% CI 51.1-78.1%) and RDD in 58.2% of the results (95% CI 44.4-71.9%) (p<0.001). The median time for the ceftazidime/avibactam susceptibility result was 10.6 h with BD Phoenix™ NMIC-502 panels, while the RDD result required 24 h incubation in 92.3% of the isolates. With BD Phoenix™ NMIC-502, results were obtained in less than 8 h in a significantly higher proportion for 8 antibiotics: aztreonam, cefepime, cefixime, ceftazidime, ceftriaxone, levofloxacin, nitrofurantoin and tigecycline. In contrast, RDD obtained a higher percentage of results at 8 h for carbapenems and cephalexin.

## DISCUSSION

Our study showed a good performance of CPO-T for carbapenemase detection in *E. coli* which only produce OXA-48 carbapenemase and all isolates were reported as ertapenem resistant. A slightly lower proportion of correct carbapenemase assignment was found than previous studies, but it should be noted that the isolates tested produce only one type of carbapenemase which display low-level meropenem and ertapenem hydrolysis (3). No reasons were found to explain the misidentification of class D as class B in five of the isolates included in our study. The assignment algorithm should include susceptibility to ceftazidime/avibactam to avoid misclassification as a class B OXA-48 producer. In previous studies (4-9) and only including OXA-48-like producing *Enterobacterales*, 215 isolates were evaluated and 90.3% were correctly classified as producers of class D carbapenemase, and misclassification was found to be lower than our data (8.4% were unclassified, 0.4% were classified as class B-producing and 0.9% were considered negative).

Several studies have reported different approaches, using colorimetric tests, immunochromatographic assays or molecular techniques to detect resistance genes directly from positive blood cultures or colonies to enable empirical treatment to be adjusted as quickly as possible (15-17). In addition to these methods, several promising new rAST systems are being developed and implemented in clinical settings (18-20). However, these instruments are usually costly, which complicates its availability in many laboratories. Based on the time to results, rAST has been defined as a technology that yields results in less than 8 hours, thus providing clinicians with the final report within the same working shift or, at least, the same day (21). This may have a direct impact on patient management (22). This goal can also be achieved with routinely used traditional methods, such as disk diffusion. In our study, both RDD and the BD Phoenix™ panel provided definitive results in less than 8 hours for several antibiotics widely used in the empirical treatments of serious infections.

Compared to our reference method, BD Phoenix™ gave the highest rates of ME for imipenem and meropenem, and of VME for imipenem. Determining resistance and susceptibility for carbapenems using disk diffusion is a challenge in some cases, because colonies grew within the zones of inhibition. This fact has been described in other studies testing carbapenem resistance in KPC-producing *Enterobacter* (23). In *E*.*coli*, rates of carbapenem-heteroresistance of 25-32%, detected by disk-diffusion, have been described (24, 25). Furthermore, for OXA-48, presence of colonies within the inhibition zones in disk or gradient diffusion assays for meropenem, independently of the MIC values measured by microdilution, has been reported in *Klebsiella pneumoniae* producing this carbapenemase (26). Previous studies have shown that the outcomes of *in vitro* susceptibility testing of tigecycline can be affected by the testing method used, mainly in *Acinetobacter baumannii* but also for carbapenemase-producing enterobacterales (27). On the other hand, the discrepancies between the absence of mutations in the targets or in the regulator of the AcrAB-TolC system, as well as the lack of acquired genes in resistant isolates by disk diffusion, are remarkable. Comparative studies of resistance phenotype prediction using sequencing data had already noted low concordance, partly because a proportion of the isolates might possess uncharacterized antimicrobial resistance genes (28).

The limitations of this study are the number of isolates included and that the reference method used is not the usual one for this type of evaluation. Very few studies include such numbers of OXA-48-producing *E. coli* isolates that do not co-produce ESBL/AmpC production. In addition, the isolate collection is multicentric and highly polyclonal, and the isolates are characterized by sequencing. The reference method was chosen as it allows comparison of read times and hence time to results, which is not possible with microdilution.

In conclusion, our results demonstrate that CPO-T is a reliable tool for simultaneous detection and classification of OXA-48 in *E. coli* isolates that do not co-produce ESBL/p*AmpC* where carbapenemase production is difficult to detect without the need for additional testing. The specific strength of carbapenemase detection in routine susceptibility testing as provided by the CPO-T lies in those samples that generally receive lower priority. BD Phoenix™ also provides definitive results in less than 8 hours for at least nine antibiotics in these isolates, including some of those most frequently used in empirical treatment.

## Acknowledgments

This research received no specific grant from any funding agency in the public, commercial, or not-for-profit sectors.

## Authors contribution

Fátima Galán-Sánchez, Conceptualization, Data curation, Formal analysis, Investigation, Methodology, Validation, Writing – original draft, Writing – review and editing | Inés Portillo-Calderón, Investigation, Methodology | Manuel Rodriguez-Iglesias, Funding acquisition, Resources, Validation, Writing – review and editing | Álvaro Pascual, Funding acquisition, Resources, Validation, Writing – review and editing |Lorena López-Cerero, Conceptualization, Data curation, Formal analysis, Investigation, Methodology, Supervision, Software, Validation, Writing – review and editing

## Supplemental Material

Figure S1: Dendogram of 51 OXA-producing *Escherichia coli* constructed based on Enterobase core genome scheme

Table S1: Whole genome sequencing quality parameters

## Data sharing

Data are contained within this article and Supplementary Materials. WGS data are available in the ENA database (Reviewer link: https://dataview.ncbi.nlm.nih.gov/object/PRJNA1104240?reviewer=m1jkbifa4hjlbvbkasb7t7rqm7).

---

**Figure S1.**
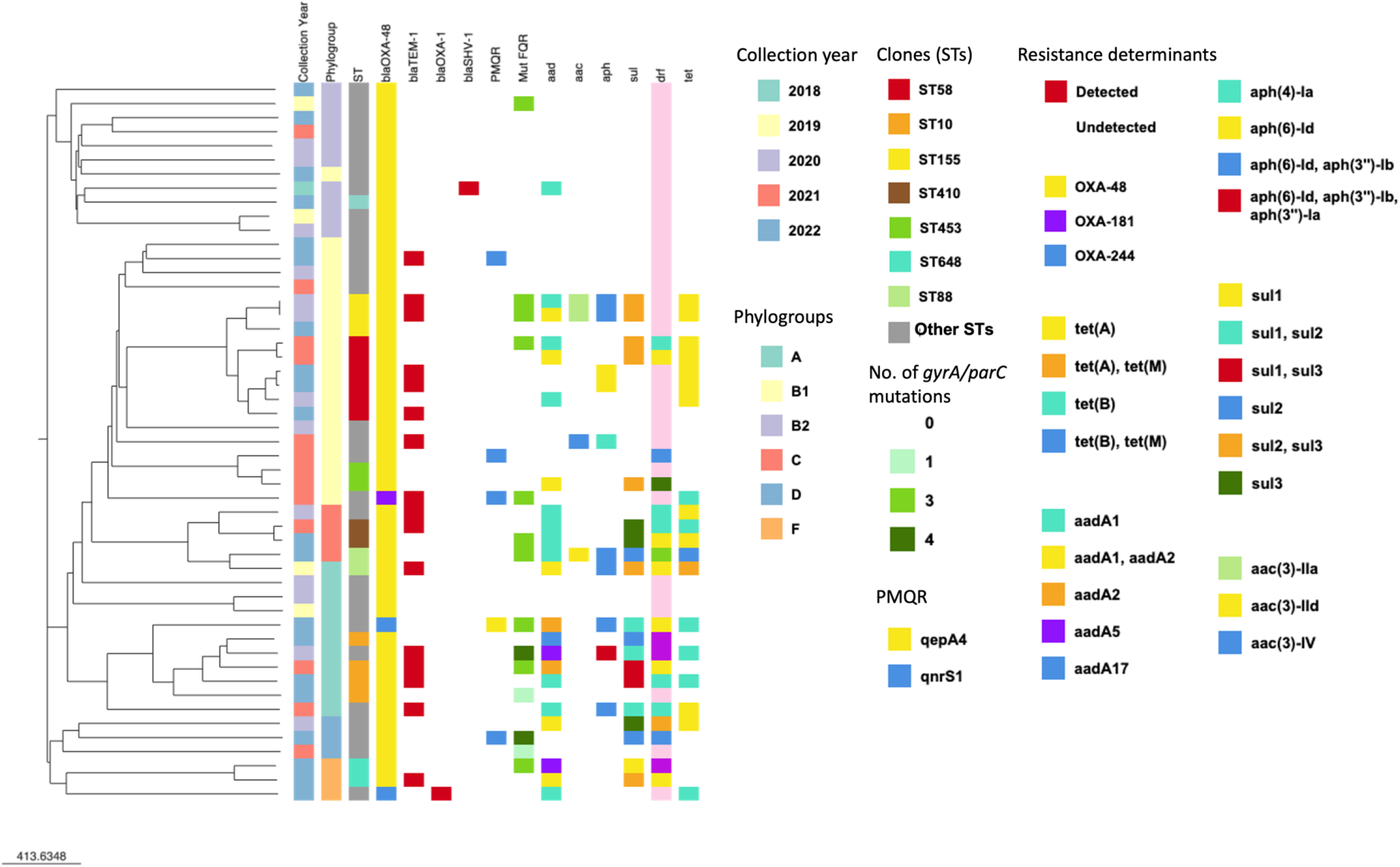
Dendogram of 51 OXA-producing *Escherichia coli* constructed based on Enterobase core genome scheme. Distribution of main resistance determinants, clones defined by MLST and phylogroups are indicated by legends colours. PMQR = plasmid-mediated quinolones resistance; Mut FQR = number of mutations in FQR region (*gyrA/parC* topoisomerases).

**Table S1.**
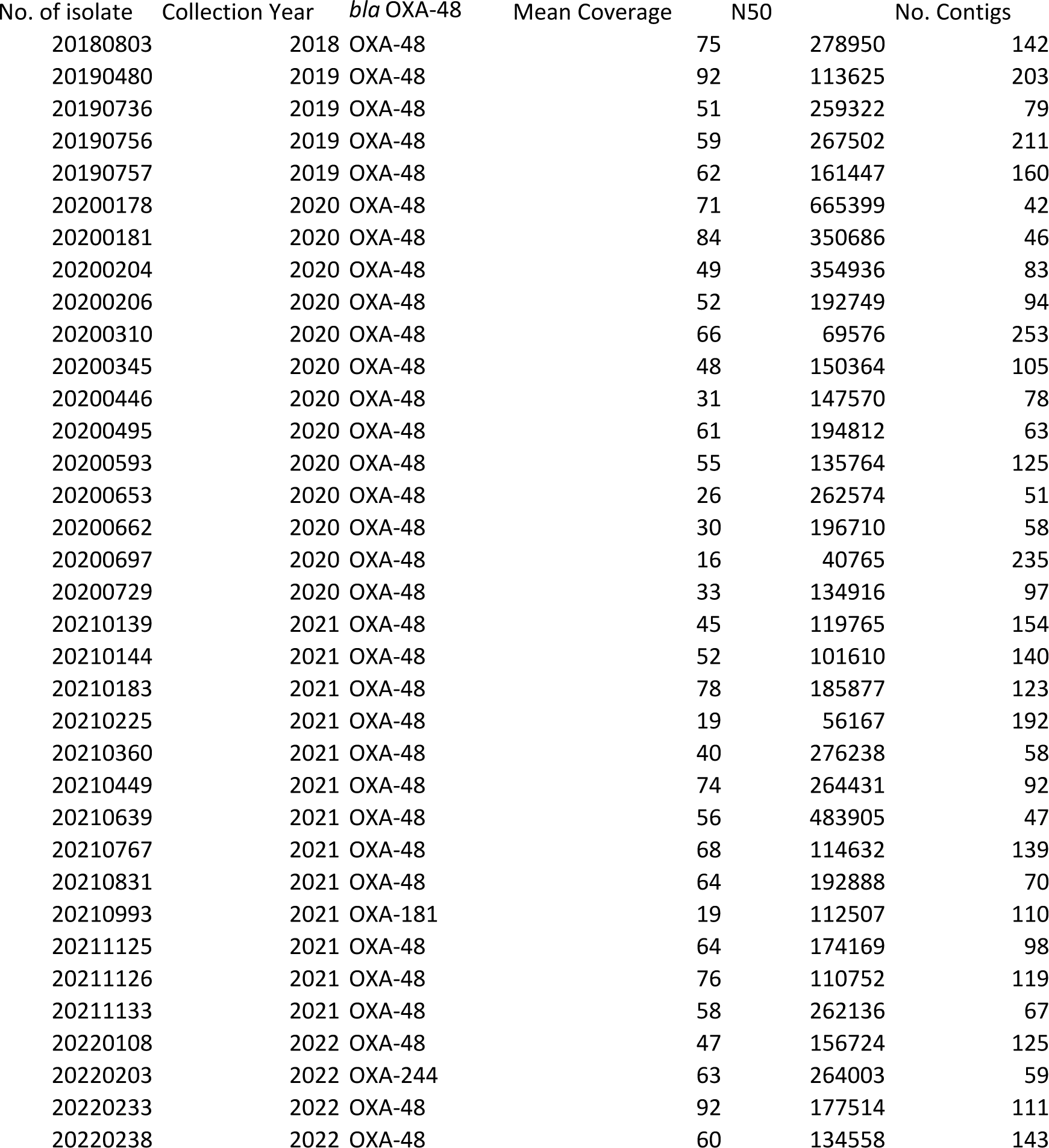
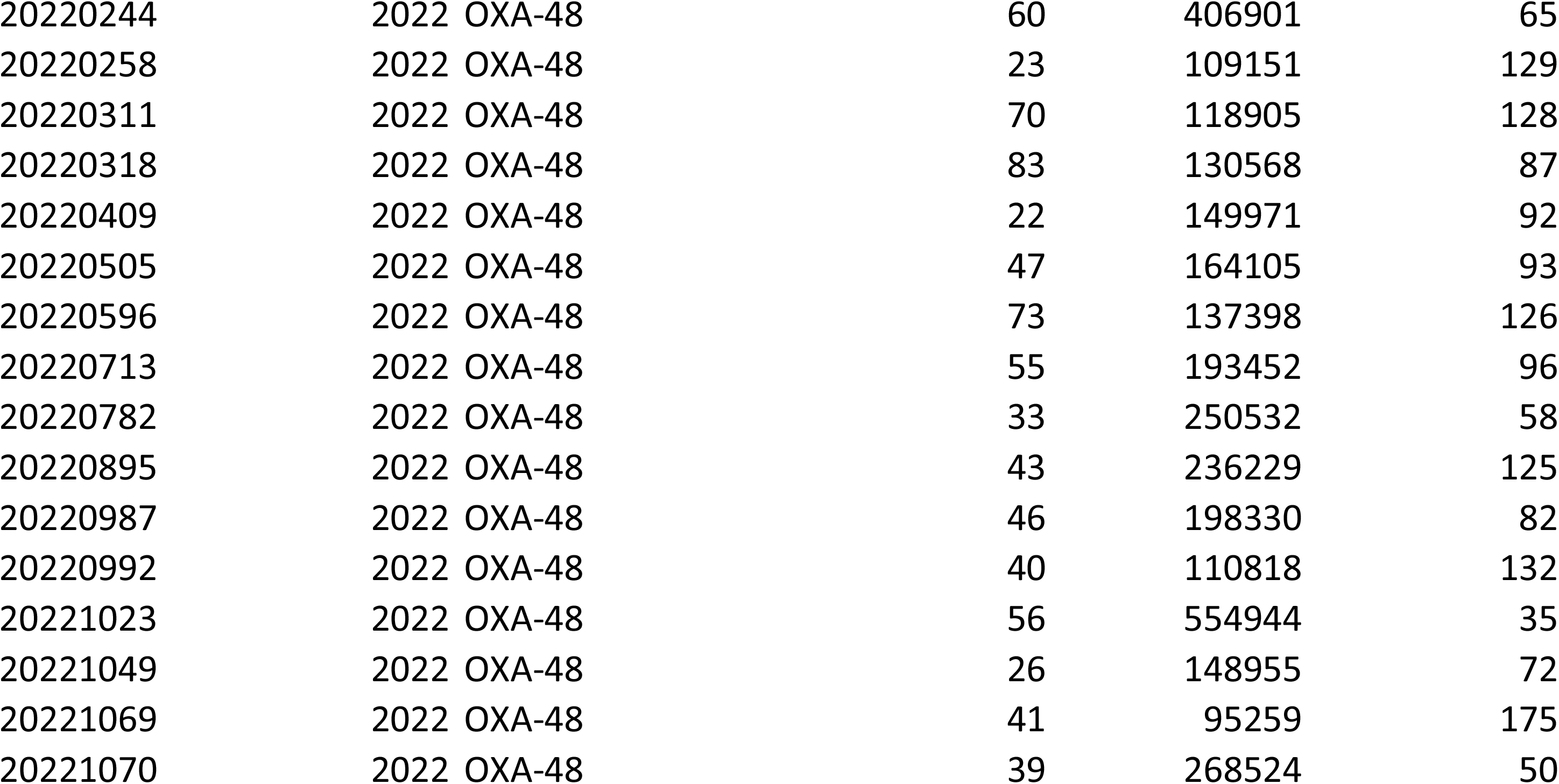
Whole genome sequencing quality parameters. N50= the sequence length of the shortest contig at 50% of the total assembly length.

